# Nanodomain formation in lipid bilayers I: Quantifying the nanoscopic miscibility transition with FRET

**DOI:** 10.1101/2025.10.28.685011

**Authors:** Emily H. Chaisson, Deeksha Mehta, Frederick A. Heberle

## Abstract

We present a robust and easy-to-use methodology for determining the nanoscopic miscibility transition temperature, nm-*T*_*mix*_, of lipid bilayer mixtures from FRET measurements. The method relies on the use of freely diffusing fluorescent donor and acceptor lipids that partition non-uniformly between coexisting phases. When this condition is met, changes in lipid clustering that occur as the sample passes through the transition result in abrupt changes in the spatial distribution of probes and consequently, abrupt changes in the FRET signal. FRET vs. temperature data can then be modeled with a phenomenological piecewise function that describes how the signal changes above and below nm-*T*_*mix*_. Using lattice simulations, we show that the transition between these regimes occurs when the size of lipid clusters surpasses a critical threshold that is approximately equal to twice the Förster distance of the donor/acceptor pair, or about 10 nm. Because other, temperature-dependent factors unrelated to lateral organization—such as changes in lipid molecular area and donor photophysics—can also influence the FRET signal, we also fit the data using a simpler model of uniform mixing. An information theory-based test comparing the fit quality of the uniform and phase separated models provides a straightforward and robust criterion for objectively assessing whether a given sample undergoes a nanoscopic miscibility transition within a temperature range of interest. We highlight the distinction between nm-*T*_*mix*_ determined by FRET (or methods with comparable spatial resolution) and the micron-scale transition temperature, μm-*T*_*mix*_, determined from diffraction-limited optical techniques. The analysis software is freely available from an online repository.

## 1. Introduction

Förster resonance energy transfer (FRET) is a photophysical process where a donor fluorophore, D, transfers its excitation energy to a nearby acceptor fluorophore, A. The probability of donor de-excitation via the FRET mechanism (as opposed to direct fluorescence emission) is high when D and A are in close contact but decreases with an inverse sixth-power dependence on the D-A separation distance. The distance at which FRET is 50% efficient is known as the Förster distance, *R*_*0*_; it is characteristic of a given D/A pair and is typically in the range of 2-10 nm. As a result, FRET is sensitive to molecular length scales, making this technique highly useful for many biological applications [1, 2].

FRET is routinely used in membrane biophysical studies to gain insight into the spatial organization of lipids and proteins both in vitro [3–8] and in vivo [9–11]. Changes in FRET efficiency can often be interpreted in terms of co-localization or segregation of fluorescently tagged species (typically lipids or proteins) under different conditions. In the context of membrane phase behavior, FRET is sensitive to abrupt changes in lipid clustering at the onset of a miscibility transition and the resulting spatial redistribution of fluorescent donor and acceptor lipids. A variety of FRET methods have therefore been devised to detect phase boundaries when composition is varied at fixed temperature [12–18], or when temperature is varied at fixed composition [19–22]. Thorough discussions of the use these methods and their relative advantages and disadvantages can be found in several comprehensive review articles [23–25].

Here, we describe a variant of steady-state probe-partitioning FRET for measuring the miscibility transition temperature (*T*_*mix*_) that requires only a single sample and access to a fluorescence spectrometer. The method exploits the abrupt changes in sensitized acceptor emission that occur as the sample passes through *T*_*mix*_. We emphasize the simultaneous use of two probe pairs with complementary partitioning behavior, which helps distinguish between genuine changes in miscibility and other factors that can trivially influence FRET efficiency, thus minimizing the possibility of misinterpretation. Using lattice simulations, we connect the observed changes in the FRET signal to a minimum cluster size of approximately 10 nm, consistent with an interpretation in terms of nanoscopic miscibility.

## 2. Materials and Methods

### 2.1 Chemicals

Phospholipids, 1,2-dioleoyl-sn-glycero-3-phosphocholine (DOPC), 1-palmitoyl-2-oleoyl-sn-glycero-3-phosphocholine (POPC), and 1,2-dipalmitoyl-sn-glycero-3-phosphocholine (DPPC) were purchased from Avanti Polar Lipids (Alabaster, AL). Cholesterol was purchased from Nu-Chek Prep (Elysian, MN). 1-palmitoyl-2-(dipyrromethene boron difluoride)undecanoyl-sn-glycero-3-phosphocholine (TFPC) was obtained from Avanti Polar Lipids, 1,1′-dioctadecyl-3,3,3′,3′-tetramethylindodicarbocyanine, 4-chlorobenzene sulfonate salt (DiD) was obtained from ThermoFisher Scientific (Waltham, MA), and naphtho[2,3-a]pyrene (naphthopyrene, Nap) was obtained from TCI America (Portland, OR). All phospholipids, dyes, and cholesterol were dissolved in HPLC-grade chloroform and stored at −20°C until use. The concentration of cholesterol (Chol) was determined gravimetrically, and the concentration of phospholipids was determined using an inorganic phosphate assay [26]. The concentration of the fluorescent dyes was determined from absorbance measurements using the following extinction coefficients: 23,800 M^−1^cm^−1^ (Nap); 96,900 M^−1^cm^−1^ (TFPC); 249,000 M^−1^cm^−1^ (DiD). Ultrapure water was obtained from a Milli-Q IC 7000 purification system (Millipore Sigma, Burlington, MA).

### 2.2 Vesicle preparation

Low lamellarity vesicles (LLVs) were prepared using the method of rapid solvent exchange (RSE) [27]. First, the desired lipid and probe mixtures were prepared by dispensing stock solutions into 13 mm x 100 mm glass screw-cap test tubes containing 25 µL of chloroform. 500 µL of ultrapure water was added to each test tube, which was then immediately mounted on the RSE device and subjected to vacuum and vigorous vortexing for 90 s. During RSE, Ar was slowly introduced into the test tube to facilitate the removal of chloroform. After 90 s, the sample was vented to Ar and transferred to a plastic centrifuge tube.

### 2.3 FRET measurements

FRET samples were prepared by combining a 0.100 mL aliquot of 0.5 mM (total lipid) LLVs with 1.900 mL of ultrapure water in a plastic centrifuge tube and vortexing before adding it to fluorometer cuvette with a flea stir bar. Fluorescence was measured using a HORIBA FluoroLog 3 spectrophotometer (HORIBA USA, Irvine, CA) equipped with a temperature-controlled cuvette holder with magnetic stirring (Quantum Northwest, Liberty Lake, WA). Each sample contained three probes comprising two D/A FRET pairs, i.e., Nap/DiD and TFPC/DiD. The probe-to-lipid ratios were 1/200 (Nap), 1/1500 (TFPC), and 1/1000 (DiD). Excitation and emission slit widths were between 2-3 nm and the signal integration time was 0.1 s. Fluorescence intensity was measured in five channels (excitation/emission wavelengths in nm): Nap direct signal (427/545); Nap/DiD FRET (427/664); TFPC direct signal (499/551); TFPC/DiD FRET (499/664); DiD direct signal (646/664). In general, the FRET channels also contain contributions from direct fluorescence emission of D and A fluorophores. To account for these contributions, signal from the FRET channel was normalized to the geometric average of the direct D and A signals using Eq. 1,

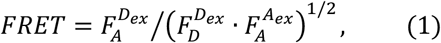

where 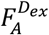 is the intensity in a FRET channel (i.e., fluorescence of the acceptor under conditions of donor excitation) and 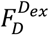 and 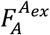 are the intensities of the corresponding direct donor and acceptor fluorescence channels, respectively [16]. To establish the temperature dependence of FRET, *FRET*(*T)* data were collected from 60 °C to 10 °C in −1 °C steps with an initial equilibration time of at least 15 min and a 2 min equilibration after each temperature change. For quantitative analysis, we used the portion of the data in the range of 15-55 °C, in part to avoid potential complications from gel phases that may be present at lower temperatures.

For each studied composition, a minimum of three samples were prepared for replicate *FRET*(*T*) measurements. Prior to analysis, FRET values were multiplied by scale factors to minimize differences among replicate datasets that result from day-to-day variability in fluorescence measurements. The scale factors for a given sample composition were determined by minimizing the sum-of-squares deviation between the FRET values of an individual replicate dataset and the mean of the replicate datasets as described in Supporting Information Section S1.

### 2.4 FRET data analysis

Because of the distance dependence of the energy transfer process, FRET efficiency is sensitive to the spatial redistribution of donor and acceptor probes that typically accompanies phase separation. In the absence of such changes—for example, when lipids remain well mixed over some range of temperature—the experimental *FRET*(*T*) data will vary gradually. In these cases, a linear function is usually sufficient for modeling small temperature ranges (< 20 °C), while larger ranges are typically better fit by a quadratic function,

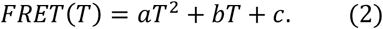

Equation 2, with its three adjustable parameters, is thus our working model for uniform mixing within the experimental temperature range. If, on the other hand, the sample passes through a miscibility phase transition as the temperature is varied, *FRET*(*T*) generally shows an abrupt change at *T*_*mix*_ A simple model that has previously been used to capture the behavior in the vicinity of *T*_*mix*_ [28] is a piecewise function of two linear regimes,

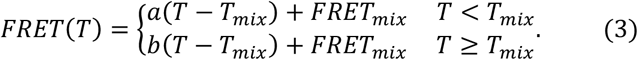

In Eq. 3, *a* and *b* are the slopes of the regimes below and above *T*_*mix*_, respectively, and (*T*_*mix*_, *FRET*_*mix*_) is the intersection point of the two regimes; all four parameters are adjusted to fit the experimental *FRET*(*T*) data.

A limitation of Eq. 3 is that FRET often exhibits non-linear behavior when the temperature is quenched well below *T*_*mix*_. A poor fit in the low-temperature regime can be avoided by restricting the analysis to data points close to *T*_*mix*_ (e.g., ± 10 °C). However, in cases where it is desirable to model a larger temperature range, an alternative approach is to substitute a quadratic function for the low-temperature regime in Eq. 3, i.e.,

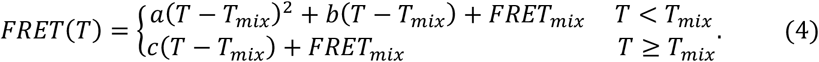

Equations 2–4 comprise a set of models of increasing complexity. Since a main objective of these experiments is to determine whether a miscibility transition exists for a particular mixture, it is typically necessary to decide whether a model for phase separation (e.g., Eq. 4) gives a substantially better fit than a uniform model (e.g., Eq. 2). When comparing two models, we are guided by the principle of parsimony: if the uniform and phase-separation models each adequately describe the data, the simpler model is favored, leading us to conclude that the membrane lipids remain uniformly mixed over the experimental temperature range at the resolution of the probe pair. To this end, we compare the corrected Akaike information criterion (AICc) of the two models to determine which provides the best description of the data [29]:

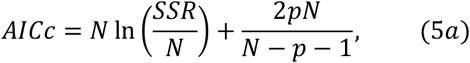

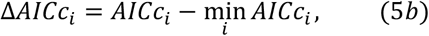

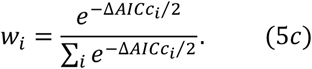

In Eqs. 5, *SSR* is the sum of squared residuals for a given model, *N* is the number of fitted data points, and *p* is the number of fitted parameters. We interpret Eq. 5c as the probability that model *i* is the most appropriate model given the experimental data and the collection of models considered.

### 2.5 Monte Carlo simulations

Monte Carlo (MC) simulations were conducted on a Mac Studio desktop computer using custom code (Mathematica v. 14.2, Wolfram Research Inc., Champaign, IL). We followed the method described by Frazier et al. [6] with some modifications. Briefly, a periodic triangular lattice of 100 x 100 sites was constructed, with each site representing one lipid molecule in a binary mixture of A and B lipids. We consider only nearest-neighbor interactions, such that the total excess lattice energy is governed solely by the unlike nearest-neighbor interaction energy, *ω*_*AB*_, and the total number of A/B contacts. Starting from an initially random configuration of A and B, lattices were equilibrated using long-range Kawasaki exchange [30], where two sites were chosen at random for a proposed exchange and the difference in energy before and after the exchange, Δ*G*, was calculated. Acceptance was determined by the Metropolis criterion [31], in which favorable or neutral moves (Δ*G* ≤ 0) are always accepted and unfavorable moves are accepted with a probability *p* = exp (−Δ*G*/*k*_*B*_*T*); for the latter, a random number was drawn from a standard uniform distribution, i.e., *X*~*U*(0,1), and the proposed exchange was accepted if *p* > *X*. Each MC cycle (*mcc*) comprised a number of proposed exchanges equal to the number of lattice sites; equilibration was judged by convergence of the total lattice energy, which required ~ 5 × 10^4^ *mcc*. All presented results are averages from 20 independent, equilibrated replica simulations.

### 2.6 Mapping the simulation lattice to Cartesian space

Estimating domain sizes and FRET efficiencies from MC simulations requires mapping the lattice site positions to a 2D Cartesian space with a nearest-neighbor distance that is appropriate for a fluid lipid membrane. This process is demonstrated graphically in Fig. S1 and briefly described here. We first define a 1D lattice site index, *k* ∈ [1,2, …, *L*^2^], where the integer *L* is equal to the number of lattice sites along the horizontal and vertical directions of a rectangular region and *L*^2^ is the total number of sites contained in the region. We also define a 2D lattice index coordinate, (*i*_*k*_, *j*_*k*_), where the integers *i*_*k*_, *j*_*k*_ ∈ [0,1, …, *L* − 1] correspond respectively to the rows and columns of lattice sites within the rectangular region and

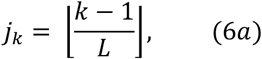

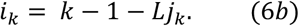

The 2D Cartesian coordinate for lattice site *k*, (*x*_*k*_, *y*_*k*_), is then calculated from the lattice index coordinate,

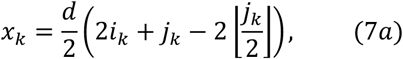

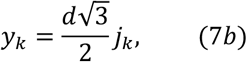

where *d* is the nearest-neighbor distance as shown in Fig. S1. To determine an appropriate value for *d*, we equate the area of a hexagonal lattice site, 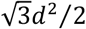, to the area per lipid, *APL*; solving for *d* yields

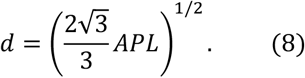

For all subsequent calculations, we used a value of 0.6 nm^2^ for *APL*, corresponding to *d* = 0.83 nm.

### 2.7 Calculating domain size from a simulation snapshot

To determine the effective cluster/domain size of a simulation snapshot, we first calculated the spatial correlation function, *g*(*r*), by applying the Wiener-Khinchin theorem [32], which states that the autocorrelation *G* of an image *I* is equal to the inverse Fourier transform of the image power spectrum *P*:

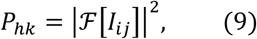

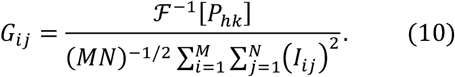

The denominator of Eq. 10 is a normalization factor, with the indices *i* and *j* corresponding to the rows and columns of pixels in the real space image. To make use of the theorem, we first converted an equilibrated lattice snapshot to a 500-pixel × 435-pixel image consisting of filled black and white hexagons corresponding to the A and B lipids. After trimming 10 pixels from each edge, Eqs. 9–10 were applied to the image to calculate *G*. Using the Cartesian pixel coordinates (see Section 2.6), *g*(*r*) was then calculated via azimuthal integration of *G*.

By definition, compositional correlations are completely lost when the correlation function decays to zero. We thus calculated the effective cluster size, *c*^∗^, as the distance where *g*(*r*) first crosses a threshold value near zero. A positive, non-zero threshold is needed because the correlation function asymptotically approaches zero from above when the system is above its critical temperature; for our simulations, this behavior is observed for *ω*_*AB*_ < 0.55 *k*_*B*_*T* [33]. We used a threshold of 0.02, as this value yielded effective cluster sizes that were similar to the domain width observed in simulations where *ω*_*AB*_ > 0.55 *k*_*B*_*T*.

### 2.8 Calculating FRET efficiency from a simulation snapshot

We calculated FRET efficiency for lattice snapshots as follows (here, we use lowercase *a* and *d* to refer to acceptor and donor species, while uppercase A and B refer to the main mixture components). In the presence of energy acceptors, the fluorescence decay of an excited state donor is given by:

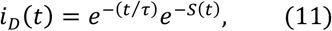

from which the energy transfer efficiency, *E*, can be calculated [34]:

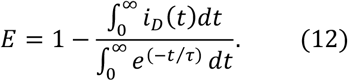

In Eqs. 11–12, *τ* is the excited state lifetime of the donor in the absence of acceptors and *S*(*t*) accounts for energy transfer arising from a particular spatial organization of acceptors. For a donor located at site *k* on a discrete lattice, *S*(*t*) can be expressed as a sum over the decay contribution from all potential acceptor sites, each weighted by the probability of finding an acceptor at that site. For a bilayer composed of two leaflets separated by a distance *h*, the energy transfer term must account for acceptors in the same leaflet as the donor as well as acceptors in the opposing leaflet, i.e., 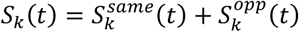, where

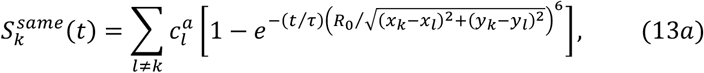

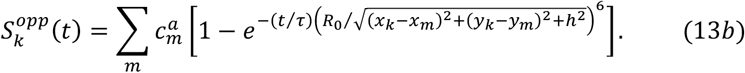

In Eqs. 13, the coefficients 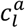 and 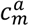 are equal to the expected number of acceptors at site *l* in the same leaflet and site *m* in the opposing leaflet, with the different indices serving as a reminder that the lattices representing the two leaflets need not be identical. These weights are thus a product of the probability of finding an acceptor at a given lattice site, 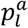, and the expected total number of acceptors on the lattice, *N*_*a*_:

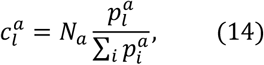

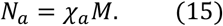

In Eq. 15, *χ*_*a*_ is the overall acceptor number density (i.e., the number of acceptors per lattice site) and *M* is the total number of lattice sites. Together, Eqs. 13–15 define the energy transfer term for donor site *k*, which together with Eqs. 11–12 serve to determine the average transfer efficiency, *E*_*k*_, experienced by that donor. The experimentally observed *E* is then the average of the transfer efficiency for donors at all lattice sites, each weighted by the probability of a donor being found at that site,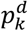:

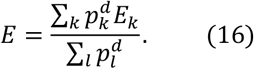

Calculating the FRET efficiency for a static 2D lattice snapshot using Eqs. 11–16 is straightforward if we know the site-by-site relative probabilities of donor and acceptor occupation, **{*p***^***d***^**}** and **{*p***^***a***^**}**. Taking inspiration from the Widom insertion method [35], we can equate these probabilities to the Boltzmann factors for inserting a probe at a particular lattice site *i*:

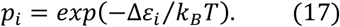

In Eq. 17, Δ*ε*_*i*_ is the difference in energy upon replacing an A or B lipid at site *i* with a probe (either *d* or *a*) and is thus determined by the set of interaction energies *ω*_*AB*_, *ω*_*pA*_, and *ω*_*pB*_ (where *p* represents either a donor or an acceptor). This treatment neglects contributions from probe-probe interactions and is thus valid in the limit of dilute probe concentration, where the probability of two probes occupying neighboring lattice sites is small.

Using the framework described above, we calculated FRET efficiency for two scenarios mimicking the complementary behavior of the two probe pairs used in our experimental studies. In both scenarios, the acceptor partitions like A, i.e., *ω*_*aA*_ = 0 and *ω*_*aB*_ = *ω*_*AB*_. In scenario 1, the donor partitions like A (*ω*_*dA*_ = 0 and *ω*_*dB*_ = *ω*_*AB*_), while in scenario 2, the donor partitions like B (*ω*_*dA*_ = *ω*_*AB*_ and *ω*_*dB*_ = 0). Scenario 1 thus mimics co-localization of *d* and *a* within A-rich domains, while scenario 2 mimics segregation of *d* and *a* between B-rich and A-rich domains, respectively. For simplicity, we used the same lattice for both leaflets of the bilayer, corresponding to perfect compositional registration. We used the following parameter values: *χ*_*a*_ = 1/400, *APL* = 60 Å^2^, *h* = 4 nm, and *R*_0_ = 3.75 nm (the same value that we calculate from the spectral overlap of Nap and DiD). In Supporting Section S2, we compare our novel method for calculating FRET efficiency for acceptors on a discrete lattice with a calculation that treats the acceptors as a continuous surface density [34]; the excellent agreement between the two methods, shown in Fig. S6, confirms the validity of our approach.

## 3. Results and Discussion

Our results are organized as follows. First, we first demonstrate the methodology for three samples with different degrees of lateral inhomogeneity: (1) DOPC, a control for a homogeneous membrane; (2) DPPC/DOPC/Chol 40/40/20 mol%, a canonical Ld+Lo mixture that exhibits micron-scale domain formation; and (3) DPPC/POPC/Chol 40/40/20 mol%, a mixture that appears uniform by fluorescence microscopy at *T* > 10 °C [36] but heterogeneous when probed with FRET [37], implying nanoscopic phase separation. We fit these datasets to models for uniform mixing (Eq. 2) and phase separation (Eq. 4), using an information theory-based metric to compare the results and thus evaluate the extent of lateral heterogeneity. For samples exhibiting phase separation, we also determine *T*_*mix*_ from model fitting and comment on the reproducibility of this measurement. We then apply the methodology to a set of samples consisting of 1:1 DPPC/DOPC and increasing cholesterol concentration to determine *T*_*mix*_ for the individual samples as well as the upper Ld+Lo phase boundary for the compositional trajectory. Finally, we use lattice simulations to correlate the value of *T*_*mix*_ obtained from *FRET(T)* data to a minimum domain size, thus informing on the spatial sensitivity of the technique.

### 3.1 Characteristics of experimental FRET data

FRET efficiency is typically quantified from donor quenching in the presence of acceptor, i.e., *E* = 1 − *F*_*DA*_/*F*_*D*_, where *F*_*D*_ and *F*_*DA*_ are the measured fluorescence intensity values of samples containing only donor or both donor and acceptor, respectively. The requirement of two samples can be circumvented by instead using sensitized acceptor emission (SAE) as the metric for FRET efficiency. The ideal SAE experiment has three criteria: (1) an excitation wavelength that excites only the donor; (2) an emission wavelength where only the acceptor emits; (3) no Rayleigh or Raman scattering at the emission wavelength. When these idealizations hold, photons counted by the detector originate entirely from the energy transfer process. In practice, although the contribution from scattering can essentially be eliminated, criteria 1 and 2 are never strictly met and the measured signal generally contains some direct fluorescence emission from both donor and acceptor. Nevertheless, these spurious contributions can be estimated (and thus accounted for) from two additional measurements in channels that primarily contain signal from direct donor and acceptor fluorescence as described in Methods. Wavelength selection for the probes used in this study is shown in more detail in Fig. S2.

To demonstrate the experimental methodology, we collected data for three compositions of known phase behavior. Figure 1 shows FRET data for DOPC (left column) and two ternary mixtures, DPPC/DOPC/Chol and DPPC/POPC/Chol (center and right columns, respectively), each at 40/40/20 mol%. In the following discussion, we refer to the Nap/DiD FRET signal as *FRET*_*ND*_ and the TFPC/DiD FRET signal as *FRET*_*TD*_; in both cases, FRET denotes the normalized sensitized acceptor emission defined in Eq. 1. These data are plotted together in the top row of Fig. 1. As mentioned previously, *FRET*_*ND*_ is expected to decrease upon Ld+Lo phase separation (i.e., when *T* < *T*_*mix*_) relative to its value in a uniformly mixed bilayer at high temperature due to segregation of donor and acceptor lipids. Conversely, *FRET*_*TD*_ should simultaneously increase due to probe co-localization within the Ld phase. We also examined the ratio of the two signals, *FRET*_*R*_ = *FRET*_*ND*_ / *FRET*_*TD*_, shown in the bottom row of Fig. 1. Combining the two signals in this way amplifies the changes that occur upon demixing and thus provides a more robust determination of *T*_*mix*_ [22]. Moreover, the simultaneous use of two probe pairs with complementary partitioning behavior helps distinguish between genuine changes in lipid mixing and other factors that can trivially influence FRET efficiency (for example, temperature-dependent changes in molecular packing that would have a similar effect on both D/A pairs) thus making the experiment less prone to misinterpretation. This is exemplified in Fig. 1 by the data for DOPC bilayers, which are known to be in a uniform, Ld phase at temperatures greater than −17 °C. Consistent with the lack of a phase transition in the observed temperature range, both *FRET*_*ND*_ and *FRET*_*TD*_ (Fig. 1a) are featureless and decrease gradually with increasing temperature, as does their ratio *FRET*_*R*_ (Fig. 1d). The modest changes are likely caused by a combination of trivial, temperature-dependent effects that do not involve major changes in lateral organization. These include the gradual increase in lipid cross-sectional area with increasing temperature [38], which increases the average separation distance between donor and acceptor chromophores, as well as temperature-dependent photophysical effects such as reduced donor excited-state lifetime and changes in probe orientational averaging. Together, these factors lead to a smooth decrease in the average FRET efficiency for both probe pairs.

**Figure 1.**
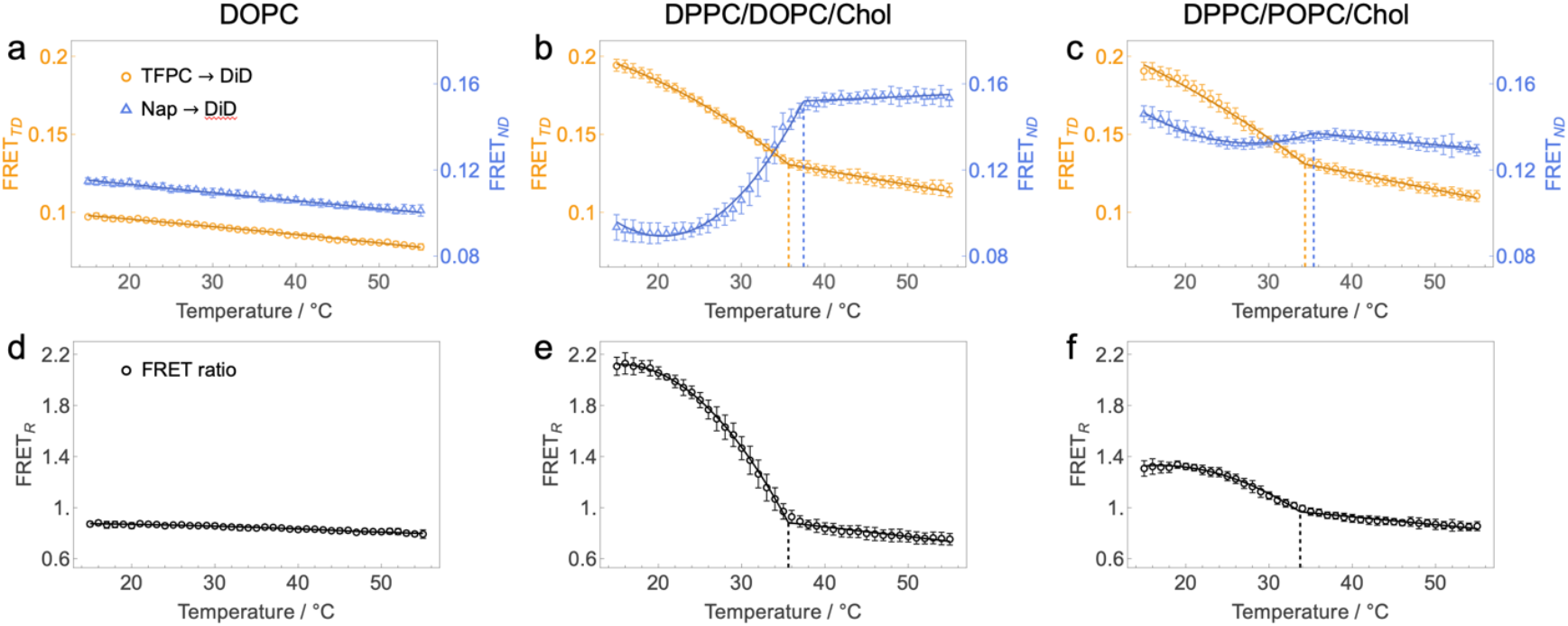
Temperature-dependent FRET reveals miscibility transitions in lipid mixtures. Shown are FRET data (open symbols) for lipid bilayers of different composition: DOPC (a, d); DPPC/DOPC/Chol 40/40/20 mol% (b, e); DPPC/POPC/Chol 40/40/20 mol% (c, f). Each data point is the average of sample replicates (*N* = 10 for DOPC and DPPC/POPC/Chol, *N* = 13 for DPPC/DOPC/Chol) and error bars are standard deviations of the replicates. The upper row (a-c) plots data for the two donor/acceptor pairs as indicated in the symbol legend, while the bottom row (d-f) plots the ratio of their FRET signals. The y-axes in a-c show the normalized sensitized acceptor emission defined in Eq. 1 rather than an absolute FRET efficiency; values are therefore dimensionless and relative. Solid lines are fits to the most appropriate model for the dataset determined from AICc as described in the text: the 3-parameter (uniform mixing) model for DOPC and the 5-parameter (phase separation) model for the ternary mixtures. The best-fit *T*_*mix*_ values are indicated by vertical dashed lines.

In contrast to DOPC, FRET data for the ternary mixtures shows a more complicated behavior at lower temperatures. As expected, *FRET*_*ND*_ for DPPC/DOPC/Chol decreases sharply when the temperature is lowered through 37 °C, while *FRET*_*TD*_ increases in slope at a slightly lower temperature (Fig. 1b). This behavior is consistent with a miscibility transition at ≈ 35-37 °C that causes a spatial reorganization of the probes, i.e., Nap partitions into the newly formed Lo domains, while TFPC and DiD remain in the Ld phase and become more concentrated as the Ld fraction decreases with decreasing temperature. Qualitatively identical trends are observed for the mixture DPPC/POPC/Chol (Fig. 1c,f), indicating that the probes partition in the same manner in nanoscopic heterogeneities as they do in optically resolvable domains. The overall magnitude of the changes is smaller for DPPC/POPC/Chol, particularly for *FRET*_*ND*_, which is likely due to weaker probe partitioning and/or smaller characteristic domain sizes in this mixture, as discussed further below.

### 3.2 Determining T_mix_ from FRET data

Although the phase behavior of the samples documented in Fig. 1 is well understood, it is desirable to have an objective criterion for determining whether a sample of unknown phase behavior exhibits a miscibility transition, as well as to precisely extract the *T*_*mix*_ value. We therefore fit each dataset in Fig. 1 to two phenomenological models: a simple 3-parameter model for the gradual variation in signal that is characteristic of uniform mixing, as well as a 5-parameter piecewise model that aims to capture the abrupt change in signal occurring at *T*_*mix*_ in the event of a miscibility transition. Unsurprisingly, goodness-of-fit for DOPC showed little improvement for the phase separation model compared to the model for uniform mixing, as demonstrated by the ratio of their sums-of-squared residuals (SSR) shown in Fig. 2a for individual replicate datasets. In contrast, for the two ternary mixtures, SSR improved substantially with the 5-parameter model, consistent with the presence of a miscibility transition in these membranes.

**Figure 2.**
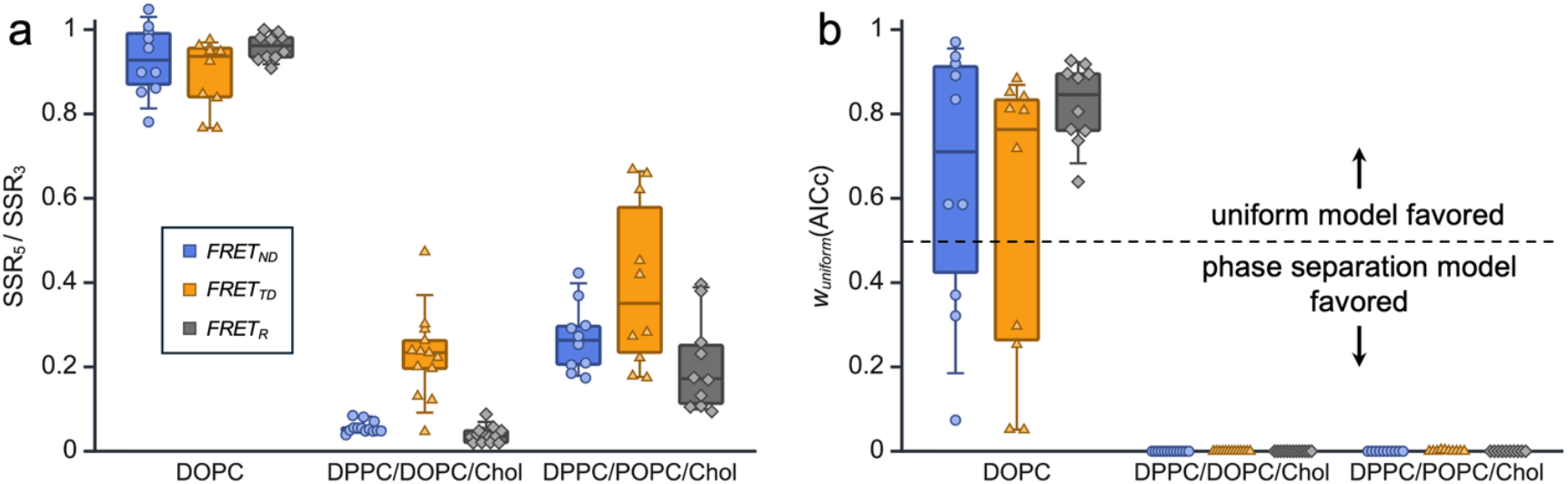
Model selection criteria for FRET analysis. (a) Box plots of the SSR ratio for the 5-parameter and 3-parameter fits (uniform and phase separation models, respectively) for each sample: from left-to-right, DOPC, DPPC/DOPC/Chol 40/40/20 mol%, and DPPC/POPC/Chol 40/40/20 mol%. Colors correspond to the two probe pairs (blue and orange) and their ratio (gray) as indicated in the legend; open symbols correspond to values from individual replicate datasets. (b) Box plots of Akaike’s weight for the uniform mixing model (categories, colors, and symbols as in panel a).

To further compare the two models, we calculated the Akaike information criterion corrected for small sample size (AICc). The difference in AICc of the 3-parameter and 5-parameter fits can be related via Eqs. 5c to the probability that the uniform model is a better description of the data; this probability is plotted in Fig. 2b for the individual probe pairs and replicate datasets. For the majority of DOPC datasets, the AICc criterion indicates that the uniform model is favored (in particular, each of the 10 *FRET*_*R*_ replicates yields a probability > 0.6), while for the two ternary mixtures, the probability shifts entirely to the phase separation model for all datasets. The most appropriate model for each full dataset (i.e., the uniform model for DOPC and the 5-parameter model for the ternary mixtures) determined by the AICc criterion is superimposed on the data in Fig. 1 as solid lines; for the ternary mixtures, the best-fit *T*_*mix*_ values are also provided in Table 1 and shown graphically as dashed vertical lines in Fig 1. It is notable that the AICc criterion indicates lateral heterogeneity in DPPC/POPC/Chol at temperatures below *T*_*mix*_ ≈ 34 °C despite the absence of visual phase separation at *T* > 10 °C when this mixture is imaged with fluorescence microscopy [36]. This result confirms that FRET is sensitive to nanoscopic heterogeneity that is otherwise undetectable with conventional optical imaging techniques. In Section 3.5, we use lattice simulations to precisely quantify the lateral resolution of these experiments.

**Table 1.** Miscibility transition temperatures from FRET data of ternary 40/40/20 mol% samples (errors are standard deviations of replicates).

| Composition | $N$ replicates | nm-$T_{mix}$ / °C | | |
| --- | --- | --- | --- | --- |
| | | $FRET_{ND}$ | $FRET_{TD}$ | $FRET_R$ |
| DPPC/DOPC/Chol | 13 | $37.3 \pm 0.9$ | $35.6 \pm 0.8$ | $35.4 \pm 0.9$ |
| DPPC/POPC/Chol | 10 | $35.2 \pm 1.7$ | $34.4 \pm 1.1$ | $34.3 \pm 0.8$ |

### 3.3 Reproducibility of T_mix_ determined from fitting FRET data

The *FRET(T)* data shown in Fig. 1 are averages of 10-13 sample replicates (the error bars correspond to the standard deviation of the replicate measurements). To evaluate the reproducibility of the *T*_*mix*_ analysis as well as highlight differences in values reported by different probe pairs, Fig. 3 plots the distributions of *T*_*mix*_ values obtained from fitting the 5-parameter model to individual replicate datasets of *FRET*_*ND*_ (blue), *FRET*_*TD*_ (orange), or *FRET*_*R*_ (gray); an example of individual replicate datasets and their fits for DPPC/DOPC/Chol is shown in Fig. S3. Notably, *T*_*mix*_ of DPPC/POPC/Chol is only 1-2 °C lower than that of DPPC/DOPC/Chol, despite the considerably higher chain melting temperature of POPC compared to DOPC (*T*_*M*_ = −2 °C and −17 °C, respectively). At higher cholesterol concentrations of 28 and 38 mol%, Pathak and London (also using FRET) reported similarly small differences in *T*_*mix*_ (< 4 °C) for DPPC/POPC/Chol mixtures compared to DPPC/DOPC/Chol [19]. These results suggest that the onset of nanoscopic heterogeneity upon lowering the temperature is mainly controlled by the chain melting temperature of the saturated lipid (*T*_*M*_ = 42 °C for DPPC).

**Figure 3.**
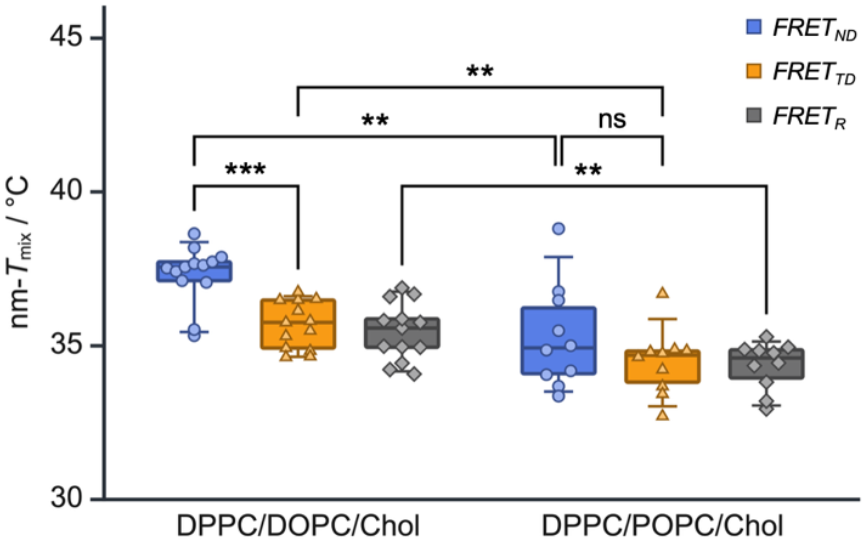
Reproducibility of miscibility transition temperatures derived from FRET. Shown are the distributions of nm-*T*_*mix*_ values obtained from sample replicates of DPPC/DOPC/Chol 40/40/20 mol% (left group, *N* = 13) and DPPC/POPC/Chol 40/40/20 mol% (right group, *N* = 10). For each replicate, *T*_*mix*_ was determined by fitting data from the individual probe pairs (blue and orange symbols for Nap/DiD and TFPC/DiD FRET, respectively) or by fitting the ratio of the signals (gray symbols) as indicated in the symbol legend. Statistical comparisons are two-tailed t-tests for independent samples with homogeneous variance: ns, not significant; *, p < 0.05; **, p < 0.01; ***, p < 0.001.

Interestingly, in both DPPC/DOPC/Chol and DPPC/POPC/Chol, *T*_*mix*_ obtained from Nap/DiD (*R*_*0*_ = 3.8 nm) is slightly higher than that obtained from TFPC/DiD (*R*_*0*_ = 4.5 nm), although the difference is not statistically significant at the 0.05 level for DPPC/POPC/Chol (Fig. 3). Using two donor/acceptor pairs with an even larger difference in *R*_*0*_, Pathak and London consistently observed a 4-6 °C difference in *T*_*mix*_ reported by NBD-DPPE/rhodamine-DOPE (*R*_*0*_ = 4.9 nm) compared to pyrene-DPPE/rhodamine-DOPE (*R*_*0*_ = 2.6 nm) in DPPC/DOPC/Chol and DPPC/POPC/Chol mixtures, with the smaller-*R*_*0*_ pair again reporting a higher *T*_*mix*_ than the larger-*R*_*0*_ pair [19]. This difference was attributed to the existence of “ultrananodomains” that could be detected only by the smaller-*R*_*0*_ pair. Our observations are consistent with this interpretation and suggest an experimental strategy for mapping how nanodomain size changes with temperature by using a set of donor/acceptor pairs covering a large range of *R*_*0*_.

### 3.4 Determining phase behavior at different cholesterol concentrations

To further demonstrate the utility of the method, we next investigated the miscibility of 1:1 DPPC/DOPC as a function of varying cholesterol concentration between 5-40 mol%. In a previous study from our group, GUVs imaged with confocal microscopy at 22 °C showed coexisting Lβ+Ld (i.e., coexisting gel and liquid-disordered) phases at and below 15 mol% Chol, coexisting Lo+Ld phases from 20-30 mol% Chol, and uniform mixing above 30 mol% Chol [22]. In the same study, we used FRET to determine that mixtures with up to 35 mol% Chol were laterally heterogeneous at temperatures below ~ 35-38 °C.

Figure 4a shows newly acquired *FRET*_*R*_ data for these compositions (symbols are replicate averages, error bars are standard deviations) overlaid with the most appropriate model fit (solid lines) as determined by the AICc criterion. Model selection is demonstrated in Fig. 4b, which plots the average and standard deviation of the AICc probability for the uniform model obtained by fitting individual replicates. Consistent with our previous analysis, the AICc criterion leaves little doubt that samples with < 35 mol% exhibit a miscibility transition between 30-40 °C. The interpretation is less clear for the 35% and 40% data, which may indicate that the size of lateral heterogeneities in these samples approaches the resolution limit of the probe pairs (discussed further below). At these high cholesterol concentrations, the membranes are most plausibly described as Lo-like, with any residual heterogeneity occurring on length scales too small to be robustly detected by the present FRET pairs.

**Figure 4.**
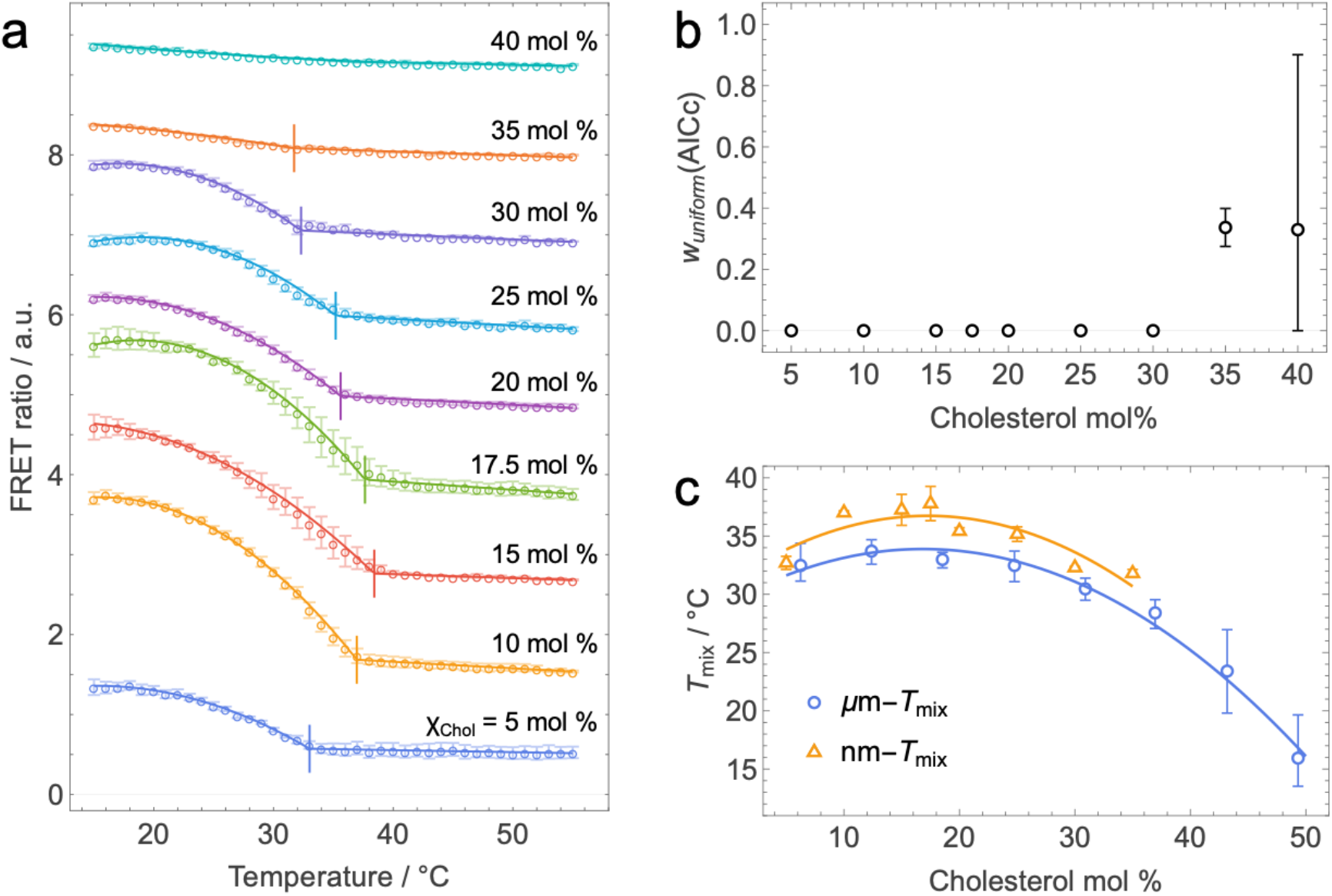
FRET data and analysis of ternary DPPC/DOPC/Chol mixtures. (a) FRET ratio vs. temperature (open circles) for liposomes composed of 1:1 DPPC/DOPC plus different amounts of cholesterol as indicated. For mixtures with < 40 mol% cholesterol, solid lines show the best fit to the 5-parameter model and short vertical bars shows the best-fit value of *T*_*mix*_. (for the 40 mol% cholesterol mixture, the solid line shows the fit to the 3-parameter model). (b) Akaike’s weight for the uniform mixing model vs. cholesterol concentration. (c) nm-*T*_*mix*_ (orange triangles) and μm-*T*_*mix*_ (blue circles, data from ref. 39) vs. cholesterol concentration. All error bars are standard deviations calculated from sample replicates.

Figure 4c plots the *T*_*mix*_ values vs. cholesterol concentration (orange triangles) determined from fitting the *FRET*_*R*_ data to the 5-parameter model; these values are listed, along with those determined from fitting the *FRET*_*ND*_ and *FRET*_*TD*_ data, in Table 2. It is notable that these values are several degrees higher than the miscibility transition temperatures reported by Veatch and Keller using fluorescence microscopy (blue circles) [39], highlighting the need to clearly distinguish between *T*_*mix*_ values determined by methods with intrinsically different spatial resolution. Here and in the companion paper [40], we refer to the miscibility transition temperature determined by confocal microscopy as μm-*T*_*mix*_ and that determined by FRET as nm-*T*_*mix*_. The sensitivity to nanoscopic domain formation is again reinforced by a small (1-2 °C) but consistent difference in the values of nm-*T*_*mix*_ reported by Nap/DiD compared to TFPC/DiD (Table 2).

**Table 2.** Miscibility transition temperatures of DPPC/DOPC 1:1 with varying cholesterol concentration (errors are SEM of replicates).

| Cholesterol mol% | $N$ replicates | nm-$T_{mix}$ / °C | | |
| --- | --- | --- | --- | --- |
| | | $FRET_{ND}$ | $FRET_{TD}$ | $FRET_R$ |
| 5 | 3 | $34.0 \pm 0.3$ | $32.7 \pm 0.7$ | $32.7 \pm 0.3$ |
| 10 | 3 | $39.4 \pm 0.1$ | $38.0 \pm 0.8$ | $37.0 \pm 0.1$ |
| 15 | 3 | $38.9 \pm 0.4$ | $37.2 \pm 0.8$ | $37.3 \pm 0.8$ |
| 17.5 | 3 | $39.6 \pm 0.7$ | $37.8 \pm 0.9$ | $37.8 \pm 0.9$ |
| 20 | 13 | $37.3 \pm 0.1$ | $35.6 \pm 0.1$ | $35.4 \pm 0.1$ |
| 25 | 3 | $36.2 \pm 0.5$ | $34.3 \pm 0.5$ | $35.2 \pm 0.4$ |
| 30 | 3 | $32.9 \pm 0.2$ | $32.31 \pm 0.01$ | $32.3 \pm 0.1$ |
| 35 | 3 | $33.5 \pm 0.4$ | $31.7 \pm 0.2$ | $31.8 \pm 0.2$ |
| 40 | 3 | $37 \pm 2$ | $35.6 \pm 0.2$ | $31.2 \pm 0.6$ |

### 3.5 Determining the characteristic length scale of nm-T_mix_

Because FRET efficiency can be exactly calculated for a given spatial arrangement of donors and acceptors, simulations are often used to gain insight into factors that influence donor and acceptor localization and thus change the FRET signal [41–46]. In the context of membrane phase behavior, Monte Carlo simulations have been successfully combined with FRET data to determine pairwise lipid interaction energies [6] and estimate domain sizes in multicomponent mixtures [6–8, 37, 47, 48].

To more precisely quantify the spatial sensitivity of our FRET method, we used Monte Carlo lattice simulations to correlate changes in FRET efficiency to cluster sizes. In these simulations, the unlike pairwise interaction energy *ω*_*AB*_ (where A and B represent the two components of a generic binary mixture) controls the extent of nonideality [49]. Figure 5a shows snapshots of equilibrated lattice configurations with increasing nonideality, starting from *ω*_*AB*_ = 0 (corresponding to ideal mixing). Small clusters are already evident at *ω*_*AB*_ = 0.2 and continue to grow up to the critical point at *ω*_*AB*_ = 0.55 k_B_T. Above this value of *ω*_*AB*_, complete phase separation occurs with the formation of A-rich and B-rich domains whose size is limited only by the dimensions of the lattice. Figure 5b shows correlation functions, *g*(*r*), computed from the lattice configurations as described in Methods. For *ω*_*AB*_ < 0.55 k_B_T, the *g*(*r*) asymptotically approach zero at large distances and are well described by the functional form *r*^−*θ*^*e*^−*r*/ *ξ*^ (i.e., the product of a power law and a decaying exponential), yielding correlation lengths, *ξ*, between 0.2 and 3 nm (Fig. S4). Because *ξ* diverges at the critical point, we also calculated an effective cluster size *c*^∗^ as the distance at which *g*(*r*) first drops below a threshold value of 0.02 (arrows in Fig. 5b). For the case of perfectly random mixing, we obtain *c*^∗^ = 0.9 nm, approximately equal to the nearest-neighbor distance. With increasing nonideality, *c*^∗^ grows gradually until the vicinity of the critical point, at which point it rapidly increases to a limiting value of ≈ 30 nm that is similar to the width of the stripe domains that form in the simulations with *ω*_*AB*_ > 0.55 k_B_T. Within this framework, changes in temperature in the experiments play a role analogous to changes in interaction strength in the simulations, primarily modulating the correlation length or effective cluster size of compositional fluctuations.

**Figure 5.**
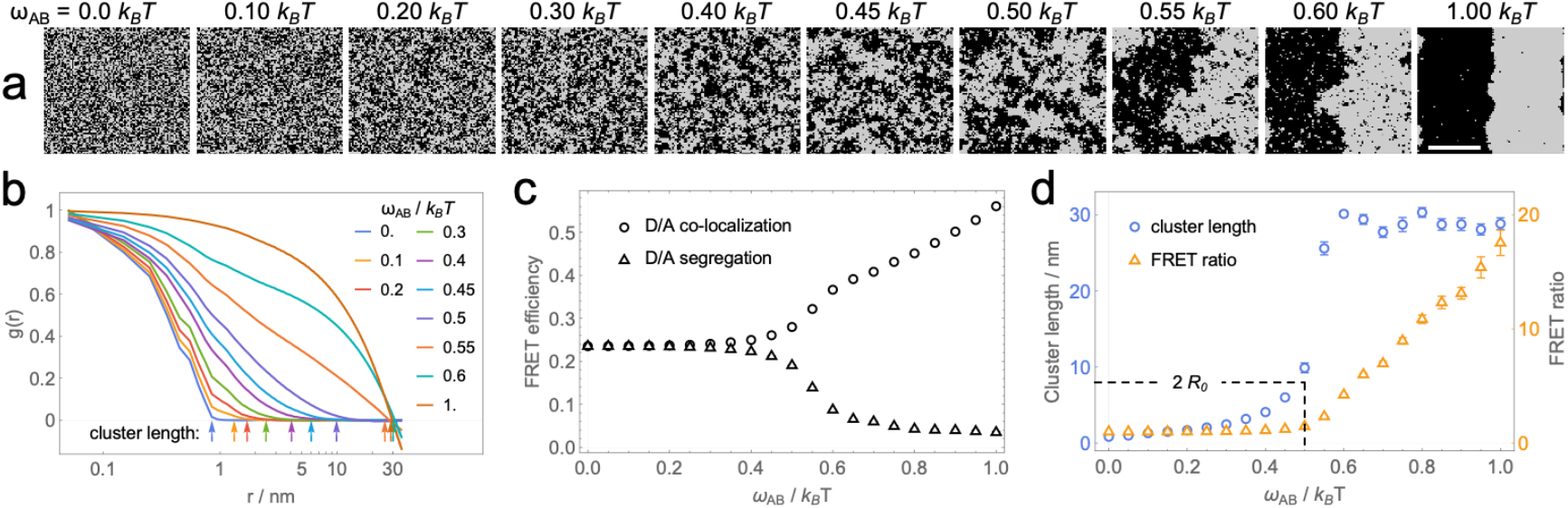
Monte Carlo simulations of lipid mixing. (a) Lattice snapshots for different values of the unlike pairwise interaction energy, *ω*_*AB*_ (scale bar is 30 nm). (b) Correlation functions computed from equilibrated lattice snapshots. The arrows mark the effective cluster size, *c*^∗^, defined as the distance at which *g*(*r*) falls below a threshold of 0.02. (c) FRET efficiency vs. *ω*_*AB*_ for donor and acceptor probes that partition into the same or different phases as indicated in the symbol legend. (d) Superimposed plots of cluster length vs. *ω*_*AB*_ and the FRET ratio vs. *ω*_*AB*_. Error bars are SEM calculated from 20 replica simulations (note that error bars in panel c are in most cases smaller than the symbol).

To relate cluster sizes to FRET signals, we calculated FRET efficiency for two scenarios corresponding to (1) colocalization of donor and acceptor within A-rich domains, or (2) segregation of donor and acceptor between A-rich and B-rich domains. Figure 5c plots FRET efficiency vs. *ω*_*AB*_ for these scenarios. These curves bear obvious similarities to the *FRET*_*ND*_ and *FRET*_*TD*_ data from ternary mixtures in Fig. 1, with increasing *ω*_*AB*_ in the simulations substituting for decreasing temperature in the experiments: just as two regimes are observed in the experimental data above and below nm-*T*_*mix*_, two regimes delineated by the critical point are seen in the simulated data. Figure 5d overlays a plot of the FRET ratio, *FRET*_*R*_, and the effective cluster size, *c*^∗^, vs. *ω*_*AB*_. Interestingly, *FRET*_*R*_ shows a distinct breakpoint at *ω*_*AB*_ = 0.5 k_B_T, at which point *c*^∗^ is approximately equal to twice the Förster distance of 3.75 nm used to calculate FRET efficiency in the simulations. This result confirms that FRET is insensitive to short-range compositional fluctuations characteristic of ideal or weakly non-ideal mixing at a temperature above the critical point, but responds strongly once the cluster size exceeds a threshold of ≈ 2 *R*_*0*_. It is important to note that this characteristic length scale depends on domain morphology: the irregular, fluctuation-dominated clusters that arise near the critical region have a large perimeter-to-area ratio, such that a larger fraction of probes are located in the heterogeneous local environment near a domain boundary. In contrast, calculations for perfectly circular domains—representing a maximally compact geometry and therefore the most sensitive case for FRET detection—show that a detectable change in FRET occurs once the domain radius exceeds ≈ 2/3 *R*_*0*_ (Supporting Section S3 and Fig. S7). Because *R*_*0*_ for most donor and acceptor lipids is in the range of 3-6 nm, a reasonable rule of thumb is that nm-*T*_*mix*_ identified by FRET marks the transition from essentially random mixing at high temperatures to a regime characterized by lateral heterogeneities that exceed 5-10 nm in spatial extent. As we show in the companion paper [40], nm-*T*_*mix*_ is generally 5-10 °C higher than μm-*T*_*mix*_ determined from conventional fluorescence microscopy, where the spatial resolution is dictated by the wavelength of visible light. This offset implies a temperature regime in which compositional heterogeneity persists on nanoscopic length scales but remains below optical resolution, consistent with Ising-like critical fluctuations.

## 4. Summary

We presented an easy-to-implement FRET method for assessing the phase behavior of a membranous sample based on a well-established information theory-based model selection criterion. The freely available code both implements the test and, when appropriate, reports the value of nm-*T*_*mix*_, thus streamlining the analysis and enabling a consistent interpretation of phase behavior. Our method exploits the simultaneous use of two probe pairs with complementary partitioning behavior (i.e., one pair that colocalizes in Ld domains and one pair that segregates between Ld and Lo) to minimize the possibility of misinterpretation. By using sensitized acceptor emission as the FRET metric, the method is amenable to single samples including cells, and the use of phenomenological models ensures that the analysis does not rely on precisely determined Förster distances and partition coefficients, nor on prior knowledge of phase fractions. Of course, this tradeoff sacrifices direct physical insight into the underlying physics of phase separation and probe partitioning that can in principle be extracted from such data. What is gained is a simple and robust methodology that delineates the regimes of uniform mixing and nanoscopic domain formation and thus facilitates consistent comparisons between different systems, from which molecular-level details can often be inferred, especially in conjunction with simulations.

We demonstrated the method by analyzing a diverse set of compositions including both uniform and phase separated mixtures, with the latter showcasing the inherent sensitivity of FRET to both macroscopic and nanoscopic domain formation. We propose the use of the terminology nm-*T*_*mix*_ and μm-*T*_*mix*_ to refer respectively to miscibility transition temperatures determined by FRET and optical microscopy. In the accompanying paper, we show that nm-*T*_*mix*_ is systematically several degrees higher than μm-*T*_*mix*_, and that the gap between these values depends on details of lipid structure [40]. We anticipate that continued exploration of the relationship between nm-*T*_*mix*_ and μm-*T*_*mix*_ will provide valuable insight into the role of different lipid species in regulating the formation and size of lipid rafts.

## Data availability

The experimental data and analysis scripts are available at https://doi.org/10.5281/zenodo.17373192.

## Author contributions

Emily Chaisson: Conceptualization, Investigation, Formal analysis, Methodology, Software, Data curation, Visualization, Writing-Original draft, Writing-Review & editing

Deeksha Mehta: Investigation, Formal analysis, Writing-Review & editing

Frederick A. Heberle: Conceptualization, Funding acquisition, Supervision, Project administration, Methodology, Software, Formal analysis, Visualization, Writing-Original draft, Writing-Review & editing

## Supporting Information

The supporting information is available for the publication at (doi) and includes 7 figures and 3 sections.

## Acknowledgements

This research was supported by NIH grant R01GM138887 (to F.A.H.).

