## Supporting Information for "Nanodomain formation in lipid bilayers I: Quantifying the nanoscopic miscibility transition with FRET"

**This PDF file includes:**

Figures S1-S7
Sections S1-S3

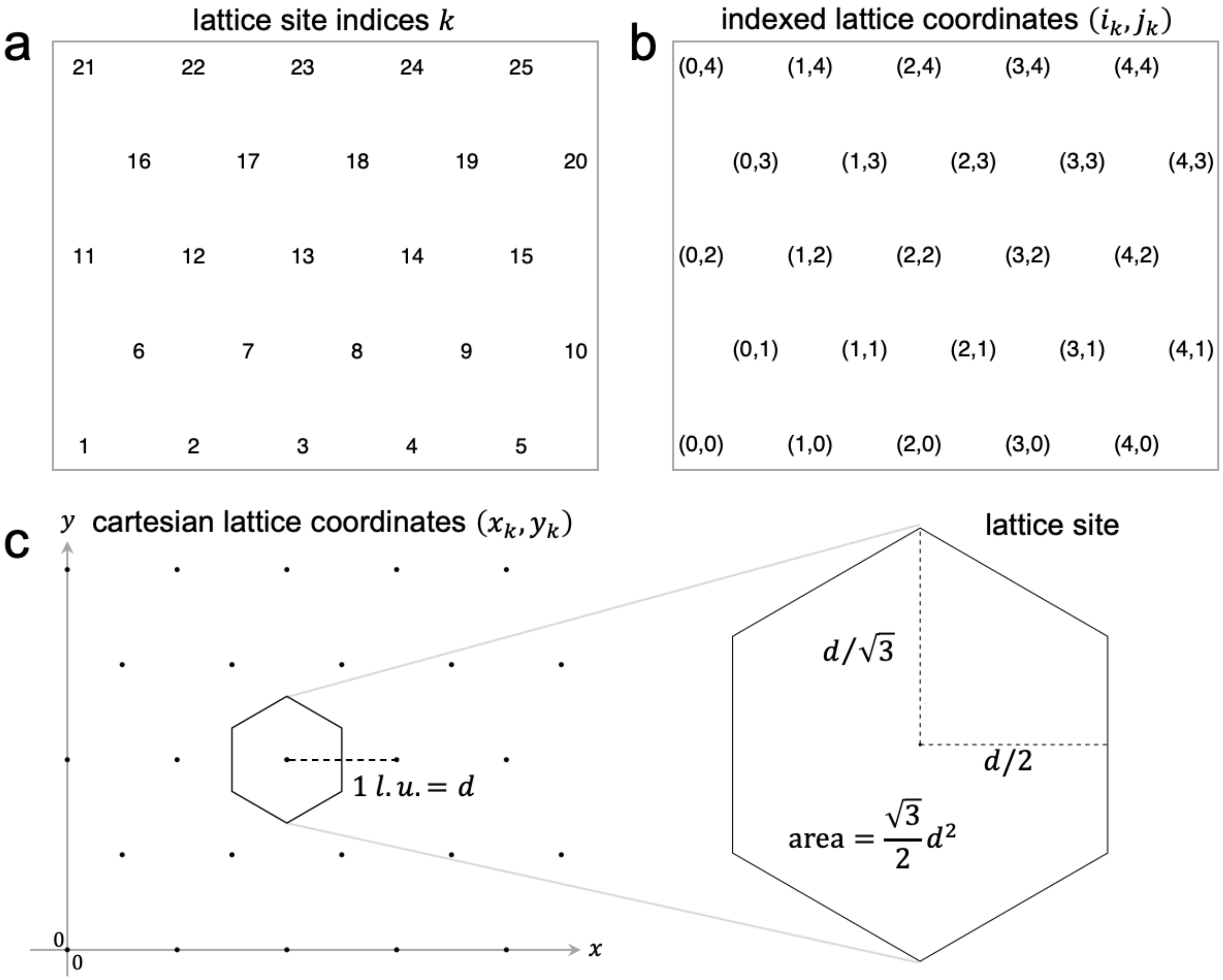


**Figure S1** Example lattice geometry for Monte Carlo simulations, here shown for a 5 x 5 lattice.

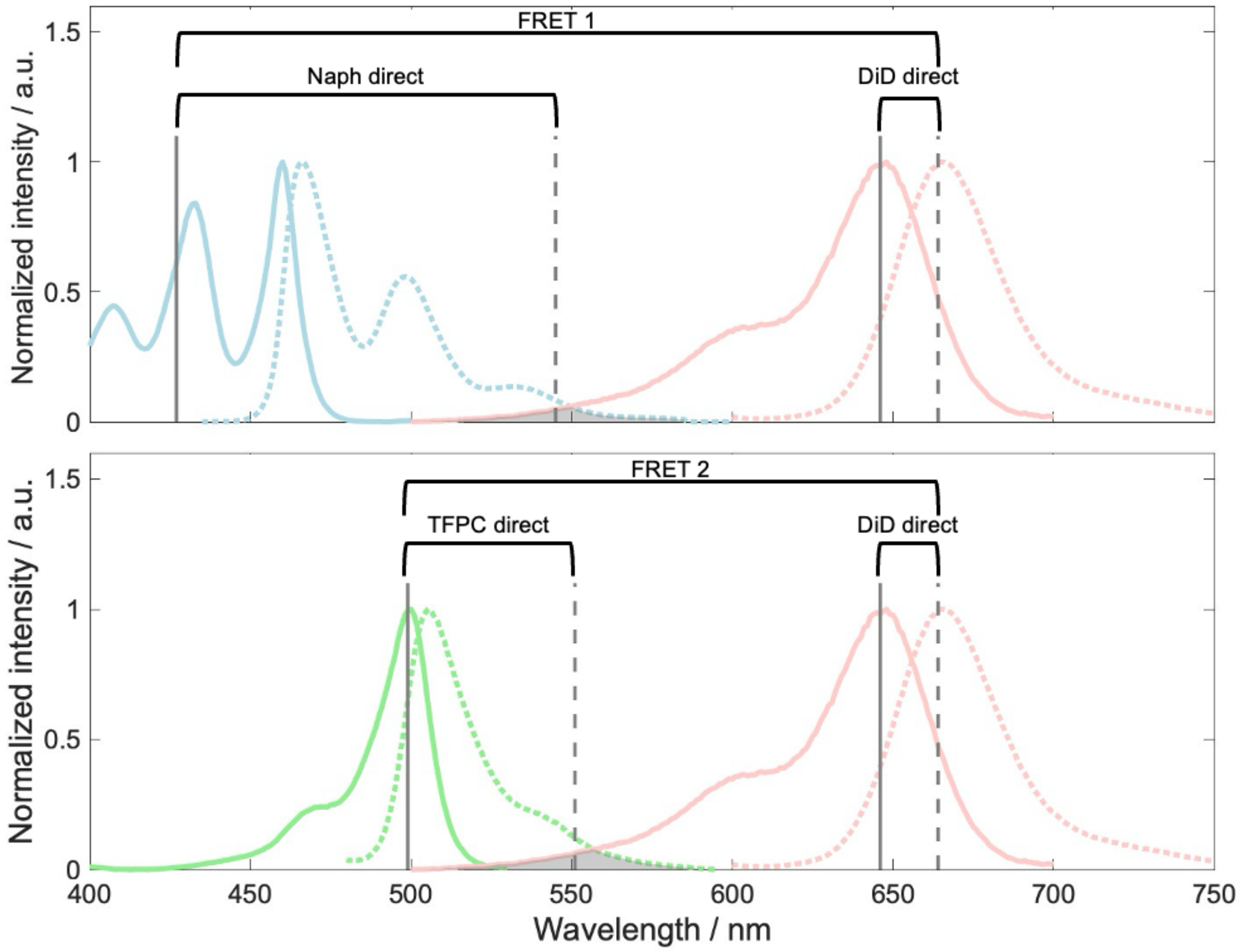


**Figure S2** Wavelength selection for probes. Solid lines are excitation spectra and dashed lines are emission spectra: Nap (blue), TFPC (green), and DiD (red). The five data collection channels (excitation and emission wavelengths) are indicated by vertical lines. The chosen excitation wavelength for Nap (427 nm) is lower than its peak excitation wavelength to avoid direct excitation of TFPC. The chosen emission wavelengths for measuring the direct fluorescence signals of Nap (545 nm) and TFPC (551 nm) are also shifted substantially from the peak emission wavelengths to avoid saturating the detector. Optimization of these wavelengths is instrument-specific and should be tailored to the particular experimental setup.

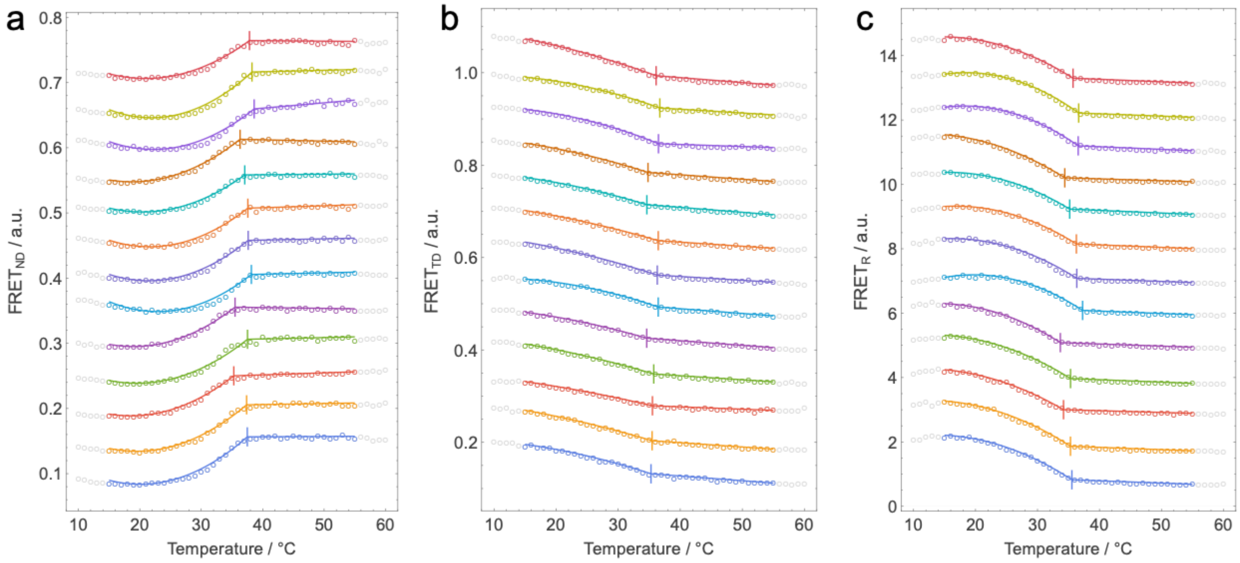


**Figure S3** Replicate FRET datasets and fits for DPPC/DOPC/Chol 40/40/20 mol%. Shown are FRET data (open symbols) and fitted curves (solid lines) for 13 replicate datasets: $FRET_{ND}$ (a), $FRET_{TD}$ (b), $FRET_{R}$ (c).

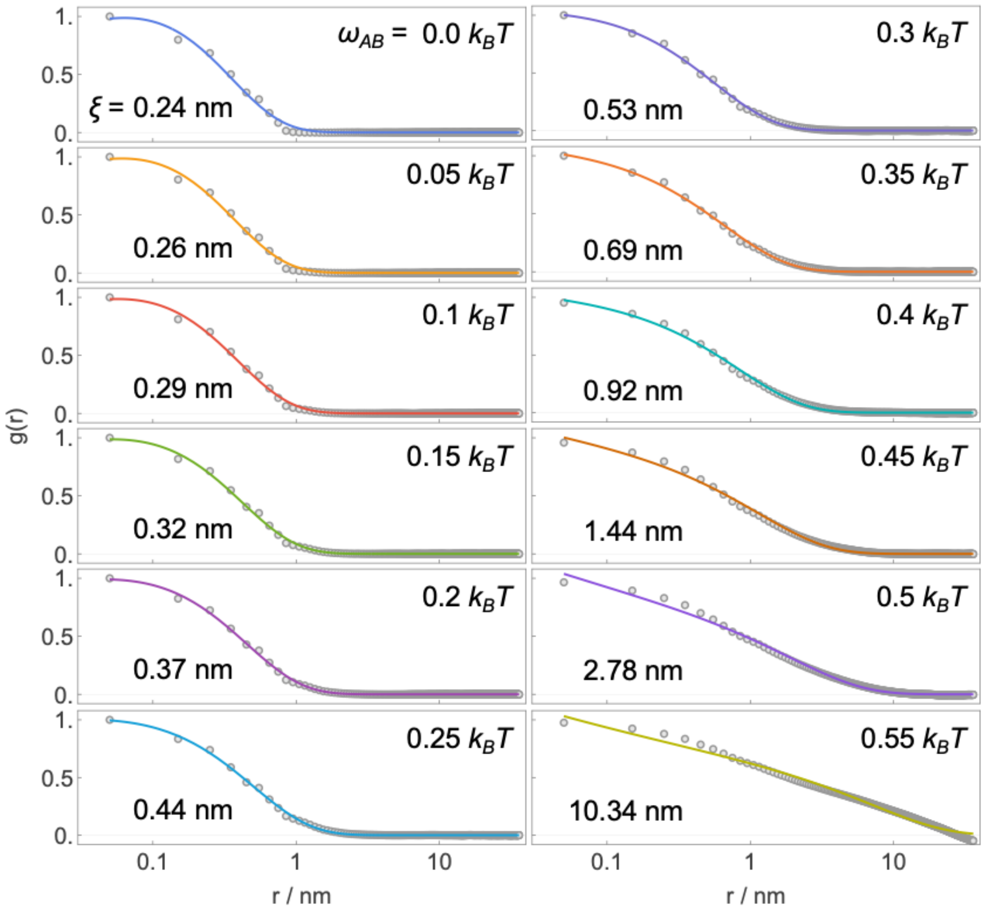


**Figure S4** Correlation lengths obtained from Monte Carlo simulations. Shown are *g(r)* data calculated from simulation snapshots (symbols) fit to the functional form $r^{-\theta}e^{-r/\xi}$ (solid lines) for values of the pairwise interaction energy $\omega_{AB} \leq 0.55$ $k_BT$. The best fit correlation lengths, $\xi$, are also shown. Each curve is the average of 20 replica simulations.

### S1. FRET data processing

Prior to quantitative analysis, FRET values were multiplied by scale factors to minimize differences among replicate datasets that result from day-to-day variability in fluorescence measurements. The scale factors were determined by minimizing the sum-of-squares deviation between the FRET values of an individual replicate dataset $f$ and the mean of the set of replicate datasets $g$ of a given sample composition and FRET pair:

$$\alpha = \frac{\sum f_i g_i}{\sum f_i^{\,2}}. \qquad (S1)$$

An example of the scaling procedure is shown in Fig. S5 for 13 replicate Nap/DiD FRET datasets from DPPC/DOPC/Chol 40/40/20 mol%.

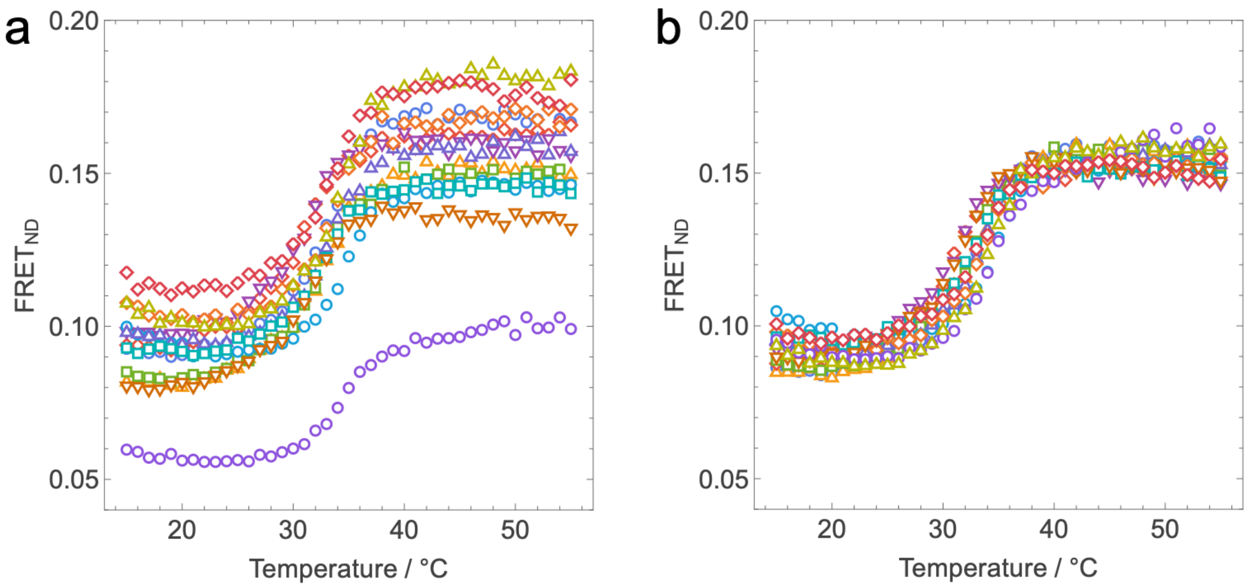


**Figure S5** Scaling procedure for replicate datasets. Shown are 13 replicate Nap/DiD FRET datasets for the composition DPPC/DOPC/Chol 40/40/20 mol%, before (a) and after (b) applying the scaling procedure described in Section S1.

### S2. FRET efficiency in a bilayer: discrete vs. continuous treatment

In this section we compare our novel method for calculating FRET efficiency for acceptors on a discrete lattice with a well-known calculation that treats the acceptors as a continuous surface density, in which case the summations in Eqs. 13 of the main text are replaced by integrals [1]:

$$S^{same}(t) = \frac{\chi_a}{APL}\int_d^\infty \left[1 - e^{-(t/\tau)(R_0/r)^6}\right]2\pi r dr, \qquad (S2a)$$

$$S^{opp}(t) = \frac{\chi_a}{APL}\int_h^\infty \left[1 - e^{-(t/\tau)(R_0/r)^6}\right]2\pi r dr, \qquad (S2b)$$

The lower limit of the integral in Eq. S2a represents the distance of closest approach between a donor and acceptor (here, equivalent to the lattice spacing $d$), while the lower limit in Eq. S2b is the distance $h$ between the two bilayer leaflets. As with Eqs. 13, Eqs. S2 are used with Eqs. 11-12 of the main text to calculate the steady-state transfer efficiency, $E$, for the bilayer [1].

Figure S6 compares $E$ vs. $\chi_a$ calculated using the continuous theory (solid lines) to that calculated using the discrete lattice treatment (open symbols) for three values of $R_0$ (2 nm, 3.75 nm, and 6 nm) and the following parameter values: $APL$ = 0.6 nm$^2$, $h$ = 4 nm, and $d$ = 0.83 nm. Because the continuous treatment corresponds to a uniformly mixed bilayer, we used lattices from the simulations with $\omega_{AB}$ = 0 (i.e., ideal mixing) for the discrete calculation. The excellent agreement between the two methods confirms the validity of our approach.

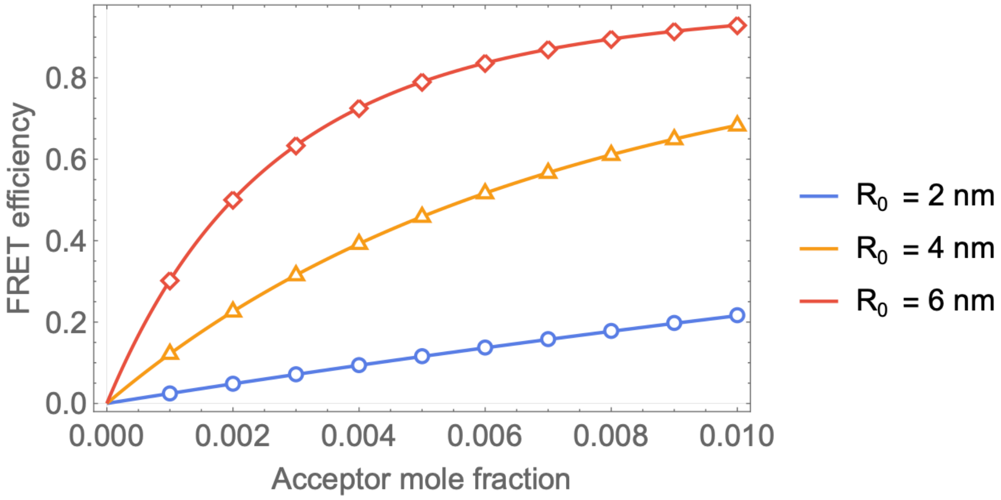


**Figure S6** Comparison of theoretical FRET efficiency to simulations. FRET efficiency vs. acceptor concentration $\chi_a$ for three values of the Förster distance, $R_0$, as indicated in the legend. Solid lines are calculated from the analytical treatment of Fung and Stryer [1] for a uniformly mixed bilayer while open symbols correspond to the average FRET efficiency calculated from replica MC lattice simulations ($N$ = 20) as described in Methods. Parameter values: $APL$ = 0.6 nm$^2$, $h$ = 4 nm, and $d$ = 0.83 nm.

### S3. FRET efficiency for circular domains

To evaluate the influence of domain shape on FRET sensitivity, we constructed lattice configurations containing perfectly circular domains. Circular domains of equal radius $R_d$ were arranged on a hexagonal lattice at an area fraction of 0.5 and superimposed onto a 100 × 100 triangular lipid lattice (Fig. S7a). The overlay was translated until the discrete fraction of lattice sites assigned to the domain phase equaled 0.5. Sites within domains were labeled as the B-rich phase and sites outside as the A-rich phase. FRET efficiency $E$ was calculated using the lattice-based framework described in the main text (Eqs. 11–16) with probe partitioning imposed via site-dependent Boltzmann weights (Eq. 17). We computed $E$ as a function of domain radius for both probe colocalization and probe segregation scenarios, and for two partitioning strengths ($K_p = 3$ and 10). In all cases, $E$ is relatively insensitive to very small domains but exhibits a clear change in slope at $R_d/R_0 \approx 0.67$ (Fig. S7b,c). Because circular domains represent a maximally compact geometry with minimal boundary length for a given area fraction, this case provides an upper bound on the sensitivity of FRET to domain formation.

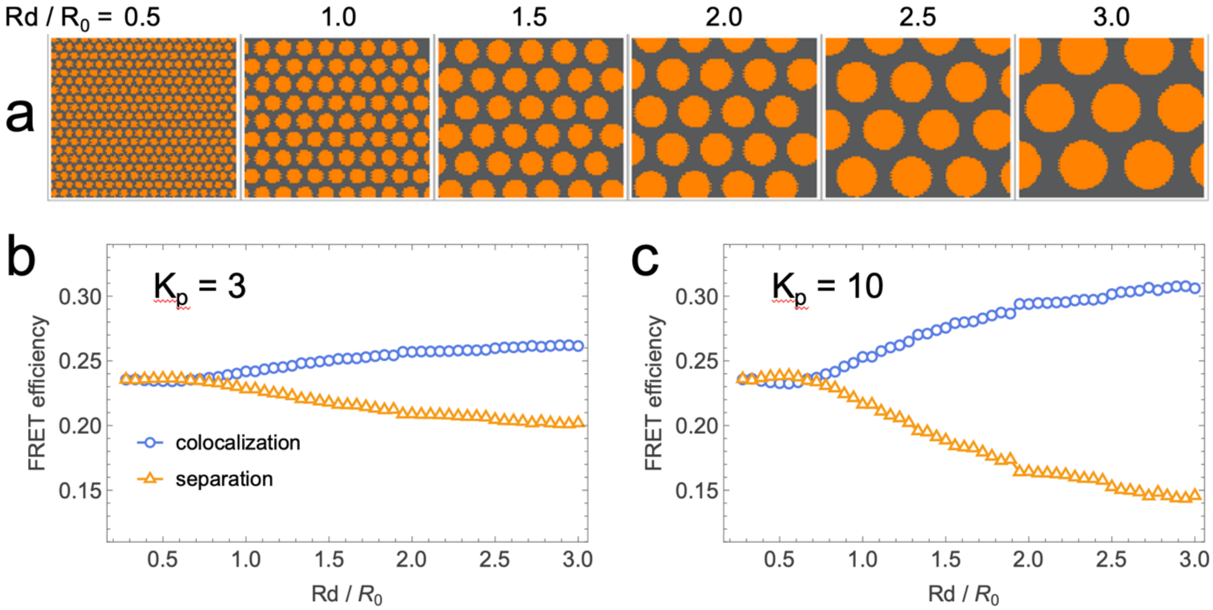


**Figure S7** Effect of circular domain geometry on FRET sensitivity. (a) Representative lattice configurations containing hexagonally packed, circular domains occupying 50% area fraction, shown for normalized domain radii $R_d/R_0$ = 0.5–3.0 as indicated. Domains (orange) are embedded in a continuous matrix phase (gray). (b,c) Calculated FRET efficiency as a function of $R_d/R_0$ for probe colocalization (blue circles) and probe segregation (orange triangles), for partition coefficients $K_p = 3$ (b) and $K_p = 10$ (c). For both partitioning strengths, FRET efficiency is weakly dependent on domain size for small domains but exhibits a change in slope once $R_d/R_0 \gtrsim 2/3$.
